# The development of ovine gastric and intestinal organoids for studying ruminant host-pathogen interactions

**DOI:** 10.1101/2021.06.30.450542

**Authors:** David Smith, Daniel R. G. Price, Alison Burrells, Marc N. Faber, Katie A. Hildersley, Cosmin Chintoan-Uta, Ambre F. Chapuis, Mark Stevens, Karen Stevenson, Stewart T. G. Burgess, Elisabeth A. Innes, Alasdair J. Nisbet, Tom N. McNeilly

## Abstract

Gastrointestinal (GI) infections in sheep have significant implications for animal health, welfare and productivity, as well as being a source of zoonotic pathogens. Interactions between pathogens and epithelial cells at the mucosal surface play a key role in determining the outcome of GI infections; however, the inaccessibility of the GI tract *in vivo* significantly limits the ability to study such interactions in detail. We therefore developed ovine epithelial organoids representing physiologically important gastric and intestinal sites of infection, specifically the abomasum (analogous to the stomach in monogastrics) and ileum. We show that both abomasal and ileal organoids form self-organising three-dimensional structures with a single epithelial layer and a central lumen that are stable in culture over serial passage. We performed RNA-seq analysis on abomasal and ileal tissue from multiple animals and on organoids across multiple passages and show the transcript profile of both abomasal and ileal organoids cultured under identical conditions are reflective of the tissue from which they were derived and that the transcript profile in organoids is stable over at least five serial passages. In addition, we demonstrate that the organoids can be successfully cryopreserved and resuscitated, allowing long-term storage of organoid lines, thereby reducing the number of animals required as a source of tissue. We also report the first published observations of a helminth infecting gastric and intestinal organoids by challenge with the sheep parasitic nematode *Teladorsagia circumcincta*, demonstrating the utility of these organoids for pathogen co-culture experiments. Finally, the polarity in the abomasal and ileal organoids can be inverted to make the apical surface directly accessible to pathogens or their products, here shown by infection of apical-out organoids with the zoonotic enteric bacterial pathogen *Salmonella enterica* serovar Typhimurium. In summary, we report a simple and reliable *in vitro* culture system for generation and maintenance of small ruminant intestinal and gastric organoids. In line with 3Rs principals, use of such organoids will reduce and replace animals in host-pathogen research.

## 1. Introduction

The mammalian gastrointestinal (GI) tract is the site of digestion and nutrient absorption, as well as a predilection site for many infectious pathogens, including bacteria, viruses and parasites. Understanding how pathogens attach and invade cells in the GI tract will help determine mechanisms of host infection, disease pathogenesis and enable strategies to prevent and control infectious disease. Both the gastric stomach and intestine share a number of common features, including a single luminal layer of epithelial cells sealed by tight junctions which is renewed approximately every 3 – 5 days. In both organs, this huge regenerative capacity is mediated by proliferation and differentiation of tissue resident adult stem cells (ASCs) (Barker et al., 2007, 2010; Sato et al., 2009; Xiao and Zhou, 2020). In intestinal tissues, pockets of leucine-rich repeat-containing G protein-coupled receptor 5 (LGR5)-expressing ASCs reside in the base of the crypts of Lieberkühn and can differentiate into all five epithelial cell types of the intestine: enterocytes, goblet cells, enteroendocrine cells, tuft cells, and Paneth cells (Barker et al., 2007; Sato et al., 2009). In the stomach the epithelia is arranged into multiple gastric units, which comprise of the gastric pit, isthmus, neck and base with proliferative stem cells located in the isthmus (Barker et al., 2010; Xiao and Zhou, 2020). The ASCs of the gastric gland can differentiate into all five epithelial cell types of the gastric stomach: surface neck mucus cells, parietal cells, chief cells, enteroendocrine cells (including G cells, D cells, and enterochromaffin-like cells) and tuft cells (Barker et al., 2010; Xiao and Zhou, 2020).

The huge regenerative capacity of GI tract and the ability of ASCs to differentiate into epithelial cell types present in the GI tract has been exploited to develop GI organoids or “mini-guts” that reflect the cellular diversity and physiology of the organ from which they were derived (Sato et al., 2009; Barker et al., 2010). Organoid models of the GI tract were first developed from mouse stomach and intestine tissues. To achieve this, researchers isolated mouse LGR5^+^ adult stem cells from these organs and cultured them in a laminin rich extracellular matrix extracted from the Engelbreth-Holm-Swarm (EHS) mouse sarcoma, with appropriate growth factors (including Wnt3a, epidermal growth factor, Noggin and R-spondin 1). The resulting organoids consisted of organ-specific tissue (gastric or intestinal epithelia) that self-organised into spherical three-dimensional (3D) structures with a single epithelial layer and a central lumen (Sato et al., 2009; Barker et al., 2010). Since this initial discovery, organoids have been derived from a large number of different tissue types and from numerous mammalian species using similar ASC isolation and tissue culture techniques.

The development of *in vitro* organoid culture systems has transformed biomedical research as they provide a reproducible cell culture system that closely represents the physiology of the host. As the majority of infectious agents enter the body or reside at mucosal surfaces, organoids derived from mucosal sites such as the gastro-intestinal, respiratory and urogenital tracts promise to transform research into host-pathogen interactions as they allow detailed studies of early infection processes that are difficult to address using animal models.

Gastrointestinal (GI) disease in small ruminants has significant implications for animal health and welfare as well as substantial economic losses because of decreased production efficiency. In sheep, gastrointestinal nematodes (GIN) have major economic and welfare impacts worldwide, with the principal GIN of sheep including: *Haemonchus contortus*; *Nematodirus battus*; *Teladorsagia circumcincta* and *Trichostrongylus* spp. (including *T. colubriformis* and *T. vitrinus*) (Nieuwhof and Bishop, 2005; Roeber et al., 2013). These parasites are transmitted by the faecal-oral route where infective stage larvae develop in either the small intestine or abomasum (which is analogous to the gastric stomach) causing significant mucosal damage associated with host inflammatory immune responses (Stear et al., 2003; Roeber et al., 2013). In addition, sheep are natural reservoirs for enteric zoonotic pathogens of worldwide significance, such as Shiga toxin producing *Escherichia coli* (STEC) and *Salmonella enterica* (Heredia and García, 2018). The obvious challenge with studying interactions between the ovine host and GI pathogens is the lack of accessibility to the site of infection, making detailed studies particularly challenging. With the current lack of physiologically relevant *in vitro* cell culture systems to study ovine-GI pathogen interactions, research has relied heavily on use of sheep infection models, which have led to important insights into host immune responses against pathogens, immune evasion by pathogens and pathogen transmission (Stear et al., 1995; McSorley et al., 2013; Ellis et al., 2014). Despite these successes, animal experiments are often complex, costly and have ethical implications.

The use of stem-cell derived GI organoids or “mini-guts” for farmed livestock species, including ruminants, is an exciting recent development that promises to provide a physiologically relevant and host-specific *in vitro* cell culture system to interrogate host-pathogen interactions (Beaumont et al., 2021; Kar et al., 2021). A recent study has demonstrated the feasibility of generating organoids from bovine ileum tissue with the derived organoids expressing genes associated with intestinal epithelia cell types (Hamilton et al., 2018). However, no ruminant gastric organoid model has been previously reported. In this current study, in line with 3Rs principles to reduce and replace the use of animals in experiments, we develop ovine ileum and abomasum organoids as physiologically relevant *in vitro* culture systems to investigate ovine GI infection and disease (Figure 1). Using RNA-seq of both tissue and derived organoids we demonstrate that the expression profile of abomasum and ileum organoids are representative of the tissue from which they were derived. In addition, we demonstrate the utility of these *in vitro* organoid systems to study host-pathogen interactions by performing challenge studies with the abomasal parasite *T. circumcincta* and enteric bacteria *Salmonella enterica* serovar Typhimurium.

**Figure 1.**
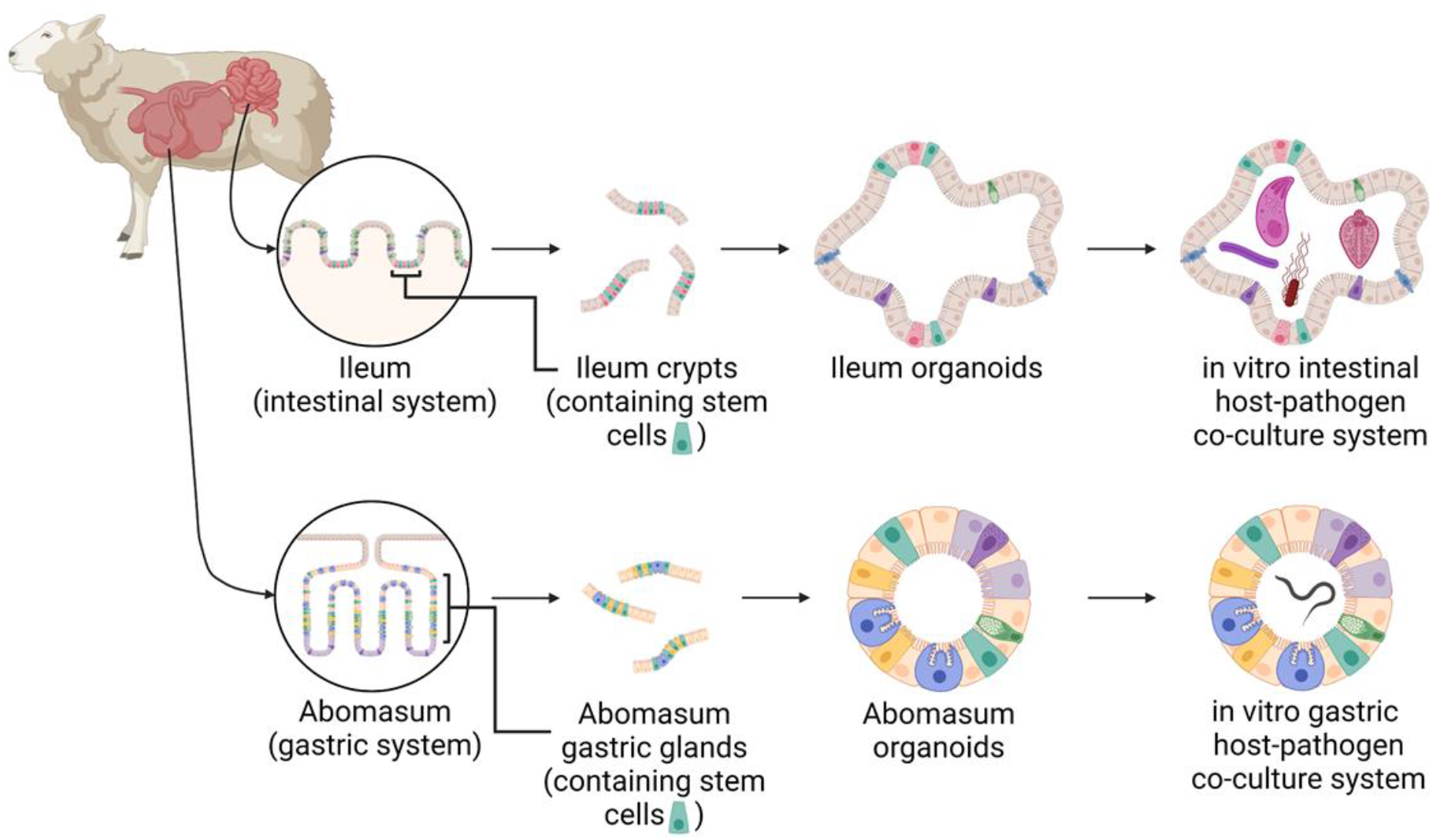
A schematic of the development of ovine gastric and intestinal organoids for studying host-pathogen interactions. Stem cells isolated from sheep ileum crypts and abomasum gastric glands can be cultivated into tissue-specific organoids when grown in a three-dimensional culture system. Gastric and intestinal organoids can be co-cultured with pathogens to model host-parasite interactions in physiologically and biologically-relevant *in vitro* culture systems. Created with https://BioRender.com.

## 2. Materials and Methods

### 2.1 Animals

All ovine abomasum and ileum tissues used in this study were derived from 7-8-month old helminth-free Texel cross male lambs (*Ovis aries*).

### 2.2 Isolation of gastric glands and intestinal crypts

Tissues were removed from sheep at post-mortem. Approximately 10 cm^2^ sections of fundic gastric fold were collected from the abomasum and approximately 10 cm sections of ileal tissue were collected from a region ~ 30 cm distal to the ileocecal junction. Tissues were removed using a sterile scalpel and forceps and placed into sterile ice-cold Hank’s buffered saline solution (HBSS) containing 25 μg/ml gentamicin (G1397-10ML; Sigma-Aldrich) and 100 U/ml penicillin/streptomycin. To expose the epithelial surfaces, the abomasum was opened along the greater curvature and the ileum opened longitudinally using dissection scissors. The luminal surfaces were rinsed with tap water to remove digesta and then placed onto sterile Petri dishes. The majority of the mucus layer was gently removed using a glass slide, after which the surface mucosal tissue (containing the gastric glands or intestinal crypts) was collected by firm scraping with a fresh glass slide. Mucosal tissue was then transferred to a Falcon tube containing 50 ml of HBSS containing 25 μg/ml gentamicin and 100 U/ml penicillin/streptomycin. Samples were centrifuged at 400 x *g* for 2 min, resulting in a tissue pellet with a mucus layer on top. The supernatant and top forming mucus layer were aspirated and discarded and the tissue was re-suspended in 50 ml of HBSS containing 25 μg/ml gentamicin and 100 U/ml penicillin/streptomycin. This process of centrifugation, aspiration and resuspension was repeated until a mucus layer was no longer visible above the pellet. To release gastric glands and intestinal crypts from tissue, pellets were re-suspended in 25 ml of digestion medium (Dulbecco’s Modified Eagle Medium [DMEM] high glucose, (11574486; Gibco) 1 % FBS, 20 μg/ml dispase (4942086001; Roche), 75 U/ml collagenase (C2674; Sigma-Aldrich) 25 μg/ml gentamicin and 100 U/ml penicillin/streptomycin) and incubated horizontally in a shaking incubator at 80 rpm for 40 minutes at 37 °C. Following digestion, the tube was gently shaken to loosen the cells and then left briefly at room temperate to allow large tissue debris to settle. The supernatant was transferred to a sterile 50 ml Falcon tube and gland/crypt integrity within the supernatant was assessed by light microscopy. Samples were then centrifuged at 400 x *g* for 2 minutes, with the resulting supernatant containing released glands or crypts. The gland/crypt-containing supernatant was washed by centrifugation at 400 x *g* for 2 minutes and the glands/crypts re-suspended in 1-2 ml advanced DMEM/F12 (12634-010; Gibco) containing 1X B27 supplement minus vitamin A (12587-010; Gibco), 25 μg/ml gentamicin and 100 U/ml penicillin/streptomycin.

### 2.4 Organoid culture

Two-hundred to one-thousand gastric glands or intestinal crypts were re-suspended in 100 μl advanced DMEM/F12 medium (containing 1X B27 supplement minus vitamin A, 25 μg/ml gentamicin and 100 U/ml penicillin/streptomycin) and were then added to 150 μl of BD Growth Factor Reduced Matrigel Matrix (356230; BD Biosciences). Fifty microliter droplets were added to consecutive wells of a 24-well tissue culture plate (3524, Corning). Plates were incubated at 37 °C, 5% CO_2_ for 15-20 minutes to allow the Matrigel to polymerize and then 550 μl of pre-warmed complete IntestiCult Growth Medium (mouse) (6005; STEMCELL Technologies) containing 500 nM Y-27632 (10005583; Cambridge Bioscience), 10 μM LY2157299 (15312; Cambridge Bioscience), 10 μM SB202190 (ALX-270-268-M001; Enzo Life Sciences) and gentamicin (50 μg/ml) were added to each well. Plates were incubated at 37 °C, 5% CO_2_ to allow organoids to develop, replacing complete IntestiCult medium every 2-3 days. Organoids were typically cultured for 7-14 days prior to passaging. Phase contrast microscopy was used to image organoids from passage one to passage five, following seven days of *in vitro* growth at each passage.

### 2.5 Organoid passage

IntestiCult media was removed from the cultured organoids and the Matrigel matrix was dissolved by replacement with 1 ml ice-cold advanced DMEM/F12. The re-suspended organoids were transferred to a 15 ml Falcon tube and the total volume of advanced DMEM/F12 was increased to 10 ml. Samples were left on ice for 5 minutes to allow organoids to settle and the supernatant was removed. The organoids were re-suspended in 200 μl advanced DMEM/F12 medium (containing 1X B27 supplement minus vitamin A, 25 μg/ml gentamicin and 100 U/ml penicillin/streptomycin) and then mechanically disrupted by repeatedly pipetting (approximately fifty times) using a 200 μl pipette tip bent at a 90° angle. The number of organoid fragments were counted by light microscopy and samples diluted to 200-1000 crypts per 100 μl. One-hundred microliters of fragments were then combined with Matrigel and plated into 24-well tissue culture plates as described in section 2.4. Phase contrast microscopy was used to image organoids from passage one to passage five, following seven days of *in vitro* growth at each passage.

### 2.6 Organoid cryopreservation

IntestiCult media was removed from the cultured organoids and the Matrigel matrix was dissolved by replacement with 1 ml ice-cold advanced DMEM/F12. The re-suspended organoids were transferred to a microcentrifuge tube and pelleted by centrifugation at 290 x *g* for 5 minutes at 4 °C. Following centrifugation, the supernatant was removed and organoid pellets were re-suspended in Cryostor CS10 cryopreservation medium (STEMCELL Technologies) at approximately 500-1000 organoids/ml before being transferred to a cryovial. Cryovials were stored in a cryogenic freezing container for 2 hours at −80 °C and subsequently transferred to −196 °C for long-term storage.

Cryopreserved organoids were resuscitated by thawing cryovials in a water bath at 37 °C and then rapidly transferring the organoids into a 15 ml Falcon tube containing 8 ml of advanced DMEM/F12 medium (containing 1X B27 supplement minus vitamin A, 25 μg/ml gentamicin and 100 U/ml penicillin/streptomycin). The cryovial was washed with a further 1 ml of media and added to the Falcon tube. Samples were pelleted by centrifugation at 290 x g for 5 minutes at 4 °C and then re-suspended in 200 μl of fresh advanced DMEM/F12 medium (containing 1X B27 supplement minus vitamin A, 25 μg/ml gentamicin and 100 U/ml penicillin/streptomycin). Re-suspended organoids were added to Matrigel and cultivated as described in Section 2.4. Organoids were imaged by phase contrast microscopy following seven days of *in vitro* growth prior to cryopreservation and post-cryopreservation.

### 2.7 Total RNA extraction

Total RNA was extracted from gastric and intestinal organoids after multiple serial passages that included passage 0 (P0) through to passage 4 (P4). Ovine gastric and intestinal organoids were prepared as described above; organoids that were formed from animal tissue-derived crypts were designated P0 and these were cultured by serial passage until P4. Each passage was cultured in triplicate wells of a 24-well tissue culture plate and allowed to mature for seven days before collecting for total RNA extraction. For total RNA extraction, IntestiCult media was removed from wells and replaced with 1 ml of ice-cold advanced DMEM/F12. The resulting suspension containing dissolved Matrigel and organoids was transferred to 15 ml sterile Falcon tubes and brought up to 10 ml with ice-cold advanced DMEM/F12. Organoids were gently pelleted by centrifugation at 200 x *g* for 5 min and the supernatant removed. Organoid pellets were re-suspended in 350 μl RLT buffer (Qiagen) containing β-mercaptoethanol, according to manufacturer’s guidelines and stored at −70°C. Total RNA was isolated from each sample using a RNeasy mini kit (Qiagen) with the optional on-column DNase digest and total RNA eluted in 30 μl nuclease-free water, according to the manufacturers protocol. Total RNA from each extraction was quantified using a NanoDropTM One spectrophotometer and integrity analysed using a Bioanalyzer (Agilent) with the RNA 6000 Nano kit. Purified total RNA was stored at −70°C until RNA-seq analysis.

Total RNA was also extracted from ovine abomasum and ileum tissue harvested at post-mortem from five individual 6-month old helminth-free Texel cross lambs and stored in RNAlater (ThermoFisher). Specifically, samples were taken from the same tissue regions stated above for crypt isolation. For total RNA isolation, approx. 30 mg of tissue was homogenized in 600 μl of RLT buffer containing β-mercaptoethanol using a Precellys® Tissue Homogenizer with CK28 tubes using x3 10s pulses at 5500 rpm with 5 min on ice between each pulse (Bertin Instruments™). Total RNA was isolated and quantified as described previously, except the total RNA was eluted in 50 μl nuclease-free water. Purified total RNA was stored at −70°C until RNA-seq analysis.

### 2.8 RNA-seq Analysis

For each sample, 1 μg of total RNA was used for RNA-seq analysis. All library synthesis and sequencing were conducted at The University of Liverpool, Centre for Genomic Research (CGR). In brief, dual-indexed, strand-specific RNA-seq libraries were constructed from submitted total RNA sample using the NEBNext® Poly(A) mRNA Magnetic Isolation Module (NEB #E7490) and NEBNext Ultra II Directional RNA Library Prep Kit for Illumina (NEB #E7760). A total of 20 libraries were constructed [including: ovine abomasum organoid P0-P4 (triplicate pooled wells for each passage); ovine ileum organoid P0-P4 (triplicate pooled wells for each passage); ovine abomasum tissues (n = 5); ovine ileum tissues (n = 5)]. The barcoded individual libraries were pooled and sequenced on a single lane of an Illumina NovaSeq flowcell using S1 chemistry (Paired-end, 2×150 bp sequencing, generating an estimated 650 million clusters per lane). Following sequencing adaptors were trimmed using Cutadapt version 1.2.1 (Martin, 2011) and reads were further trimmed using Sickle version 1.200 (Joshi and Fass, 2011) with a minimum window quality score of 20. Reads shorter than 15 bp after trimming were removed. Sequence reads were checked for quality using FastQC v0.11.7. Reads were pseudo-aligned to the *Ovis aries* transcriptome (Oar_v3.1 GCA_000298735.1) using Kallisto v0.46.2 with default settings (Bray et al., 2016) and read abundance calculated as transcripts per million (TPM). Gene expression data was analysed by principal component analysis (PCA) using pcaExplorer version 2.12.0 R/Bioconductor package (Marini and Binder, 2019). Specific genes were also manually retrieved from our transcriptomic dataset and their TPM values log2 transformed for presenting in heat maps, which were generated using GraphPad Prism software (v8.0).

### 2.9 Immunohistochemistry

Abomasum and ileum organoids were cultivated in Matrigel for 7 days in 8-well chamber slides (354118; Falcon) as described in section 2.4. To make organoids accessible to immunohistochemistry reagents, the culture medium was removed and replaced with ice-cold 4% paraformaldehyde. For fixation, samples were kept at 4 °C for 20 minutes to also dissolve the Matrigel and prevent it from re-solidifying. Organoids were washed twice with IF buffer (0.1% Tween20 in PBS) and then permeablised with 0.1% TritonX-100 in PBS for 20 minutes at room temperature. Samples were washed three times with IF buffer and then blocked for 30 minutes with 1% BSA in IF buffer at room temperature. Next, primary antibodies diluted in blocking solution were added to the organoids and samples were left overnight at 4 °C. Primary antibodies used included polyclonal rabbit α-Ki67 (ab15580, abcam, used at a 1:500 dilution), polyclonal rabbit α-EPCAM (orb10618, Biorbyt, used at a 1:600 dilution), monoclonal mouse α-villin (sc-58897, Santa Cruz Biotechnology, used at a 1:200 dilution) and monoclonal mouse α-pan cytokeratin (used at a 1:100 dilution). For isotype controls, mouse or rabbit IgG were used in place of the specific primary antibodies and were diluted at 1:100 or 1:500 for mouse and rabbit IgG respectively. The next day, samples were washed three times with IF buffer and then secondary antibodies added (diluted at 1:500 in blocking buffer) and incubated at room temperature for 1 hour. Secondary antibodies used were goat α-mouse Alexa Fluor 488 (ab150117, abcam) and goat α-rabbit Alexa Fluor 488 (ab150081, abcam). Phalloidin-iFluor 555 reagent (ab176756, abcam, used at a 1:1000 dilution) was also added during the secondary antibody step to label F-actin. Samples were washed three times with IF buffer and then Hoechst 33258 solution diluted 1:200 in IF buffer was added to label nuclei (94403, Sigma-Aldrich). Samples were incubated for a further 5 minutes at room temperature before three washes with IF buffer. Finally, slides were mounted using ProLong Gold antifade mountant (P10144, ThermoFisher Scientific) and imaged by confocal microscopy using a Zeiss LSM 710 Inverted Confocal Microscope and Zeiss Zen Black operating software.

### 2.10 Exsheathment of *Teladorsagia circumcincta* third stage larvae (L3)

*T. circumcincta* L3 (Moredun isolate MTci2, CVL) were exsheathed and labelled using modified protocols previously published (Dinh et al., 2014; Bekelaar et al., 2019). Nine milliliters of Earle’s balanced salts solution (EBSS) buffer in a 15 ml Falcon tube was preheated in a water bath to 37 °C and CO2-saturated over 1 hour using an incubator tube connected to a CO_2_ tank. Approximately 5×10^4^ *T. circumcincta* L3 in 1 ml of tap water were added to the CO_2_-saturated EBSS and the sample continued to be saturated for a further 15 minutes. The Falcon tube was then sealed with Parafilm® M and inverted 6 times before being placed horizontally into an incubator at 37 °C, 5% CO_2_ for 4 hours. Following incubation, the whole sample was transferred into a 25 cm^2^ vented cap flask and incubated overnight at 37 °C /5% CO2, to allow L3s to continue exsheathing. Exsheathment was validated the following morning by light microscopy. The larvae were then washed 4 times by repeated centrifugation at 330 x *g* for 2 minutes and re-suspension in 50 ml of distilled water (pre-warmed to 37°C). After the final wash, the L3 larvae were re-suspended in 1 ml distilled water and transferred to a microcentrifuge tube. Exsheathed L3 (exL3) were fluorescently labelled by the addition of 2 μl PKH26 dye (1 mM stock concentration) from the MINI26 PKH26 Red Fluorescent Cell Linker Kit (Sigma-Aldrich) and mixed by pipetting. Parasites were incubated with the dye for 15 minutes at room temperature, protected from light. Excess dye was removed by washing the larvae five times with distilled water as described above before finally re-suspending them in 1 ml of complete IntestiCult organoid growth medium.

### 2.11 Teladorsagia circumcincta L3-organoid co-culture

Abomasum and ileum organoids were cultivated in Matrigel for 7 days in 8-well chamber slides (354118; Falcon) as described in section 2.4. Immediately prior to organoid-*T. circumcincta* co-culture, complete IntestiCult media was removed from the cultured organoids and replaced with 250 μl of fresh pre-warmed complete IntestiCult. Twenty to 50 PKH26 labelled *T. circumcincta* exL3 in 50 μl complete IntestiCult media were added to each well of organoids and organoid-larval cultures incubated at 37°C, 5% CO_2_. Note that organoids were not removed from their Matrigel domes prior to the addition of *T. circumcincta* L3. Upon observation of multiple organoids containing *T. circumcincta* L3 within their lumen (after ~24-48 hours of organoid-*T. circumcincta* co-culture) the samples were fixed with 4% PFA for 30 min, followed by 3 washes with PBS, and stored at 4°C until fluorescence staining. Organoids were permeabilized, blocked and probed with Phalloidin-488 and Hoechst 33258 as described for organoid immunohistochemistry above. Images were captured using a Zeiss LSM 710 Inverted Confocal Microscope and Zeiss Zen Black operating software.

### 2.12 Generation of apical-out organoids

Epithelial polarity was inverted in gastric and intestinal ovine organoids by following a previously published method for reverse polarity in human intestinal organoids (Co et al., 2019). Briefly, gastric and intestinal organoids were grown in Matrigel as described above for 7 days. Matrigel domes containing developed organoids were gently dissolved by the addition of 500 μl ice-cold 5 mM EDTA in PBS, taking care not to rupture the organoids. The resulting suspension was transferred to a 15 ml Falcon tube that was subsequently filled with 14 ml of 5 mM EDTA in PBS. Samples were placed on a rocker and mixed gently for 1 hour at 4 °C. Organoids were pelleted by centrifugation at 200 x *g* for 3 min at 4 °C and the supernatant was removed. Pellets were re-suspended in complete IntestiCult growth media (containing 500 nM Y-27632, 10 μM LY2157299, 10 μM SB202190 and gentamicin (50 μg/ml), with the addition of 10 % advanced DMEM/F12 medium (containing 1X B27 supplement minus vitamin A, 25 μg/ml gentamicin and 100 U/ml penicillin/streptomycin). Re-suspended organoids were transferred to the wells of 8-well glass chamber slides and incubated at 37 °C, 5 % CO2 for a period of 72 hours, prior to being fixed and stained with Phalloidin-iFluor 555 reagent and Hoechst 33258, as described in section 2.9. Confocal imaging was performed as described in section 2.9.

### 2.13 Infection of apical-out organoids with *Salmonella enterica* serovar Typhimurium

The polarity of gastric and intestinal organoids was inverted as described above. *Salmonella* Typhimurium strain ST4/74 was chosen for this experiment as its full genome sequence is available (Richardson et al., 2011) and it has been shown to efficiently invade the ovine ileal mucosa and elicit inflammatory responses in an ovine ligated ileal loop model (Uzzau et al., 2001). To aid visualization of the bacteria in organoids, the strain was electroporated with plasmid pFPV25.1 which carries *gfpmut3A* under the control of the rpsM promoter resulting in the constitutive synthesis of green fluorescent protein (Valdivia and Falkow, 1996). Stability of the plasmid in the absence of antibiotic selection during *Salmonella* infection has been confirmed (Vohra et al., 2019). The bacteria were grown on Luria Bertani (LB) agar supplemented with 100 μg/ml ampicillin at 37 ºC overnight. Single colonies were transferred to LB broth supplemented with the same antibiotic and grown for 20 hours shaking at 180 rpm at 37 ºC. The liquid cultures were diluted to 3.3 x 10^6^ CFU/ml in complete IntestiCult growth medium, described above, and 300 μl of the dilution was added to half of the wells, which already contained organoids that had already been maintained in conditions for generating apical-out organoids for 72 hours. The other half of the wells acted as negative controls, with organoids being re-suspended in 300 μl of complete IntestiCult growth medium alone (no bacteria). After 30 minutes of incubation another 300 μl of complete IntestiCult growth medium with 200 μg/ml gentamycin was added to kill extracellular bacteria. The slides were incubated at 37 ºC, 5% CO_2_ for a total of 6 hours. At the end of the incubation period the entire volume of the liquid from each well, including the organoids, were transferred to separate 15ml Falcon tubes (Corning, UK). All centrifugations for organoid collection during washing were done at 200 rpm for 5 minutes. The supernatant was removed and the organoids were washed twice in PBS, and then re-suspended in 4% PFA for 30 minutes for fixation. The organoids were processed for immunohistochemistry as described in section 2.9 and stained with Phalloidin-iFluor 555 reagent, prior to mounting with ProLong Diamond antifade mountant (P36961, ThermoFisher Scientific). Confocal imaging was performed as described in section 2.9.

## 3. Results

### 3.1 Growth of ovine gastrointestinal organoids *in vitro*

Fragmented gastric glands and intestinal crypts isolated from the abomasum fundic fold and the ileum of 7 to 8-month old Texel cross lambs were embedded in Matrigel and grown in complete IntestiCult organoid growth medium. Under identical growth conditions, epithelial stem cells from these two different organ tissues were able to develop into organoids *in vitro* (Figure 2A). By 24 hours, sealed spherical structures containing a central lumen had formed in both the abomasum and ileum organoids. However, while the ileum organoids became branched after 5-7 days of *in vitro* culture, the vast majority of abomasum organoids retained a spherical structure that persisted for the duration of a culture passage (Figure 2A, B).

**Figure 2.**
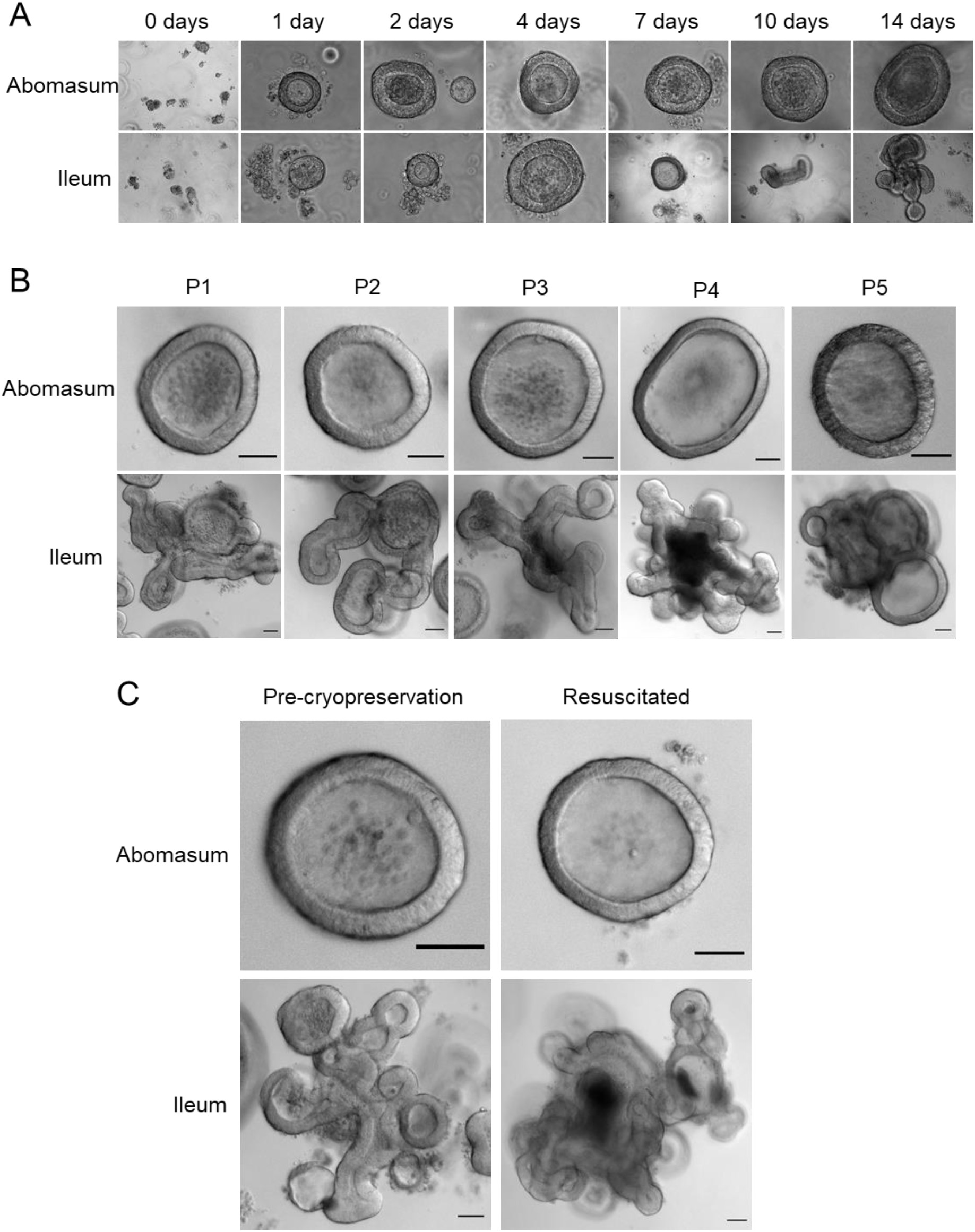
*In vitro* growth of ovine abomasum and ileum organoids. (A) Representative images of abomasum and ileum organoids grown over 14 days in the same culture conditions. (B) Representative images showing the growth and development of mature abomasum and ileum organoids across multiple consecutive passages (P1 - P5). (C) Representative images of abomasum and ileum organoids grown for seven days, both pre-cryopreservation and after resuscitation. Scale bars = 10 μm.

Abomasum and ileum organoids could be serially passaged by removal from Matrigel, fragmentation by pipetting and re-embedding in Matrigel. At each passage, ileum organoids continued to form into branched structures, while the abomasum organoids persistently formed spherical structures. (Figure 2B). Organoids that were cryopreserved in liquid nitrogen after 7 days of *in vitro* culture could be thawed and re-cultured, demonstrating the potential to store organoids long-term and to resuscitate when required. Furthermore, we found that the cryopreserved organoids can be resuscitated after at least 18 months of storage in liquid nitrogen. Abomasum and ileum organoids retained their spherical and branched structures, respectively, following resuscitation and 7 days of *in vitro* culture (Figure 2C).

### 3.2 Epithelial cell markers associated with ovine gastric and intestinal organoids

Immunohistochemistry was performed to identify key structural features associated with both abomasum and ileum organoids. Individual Z-stack images of organoids stained with phalloidin to label F-actin clearly demonstrated that the apical surface of the epithelium is present on the interior of the organoid, for both abomasum and ileum organoids, indicated by the presence of a solid F-actin-positive boundary (Figure 3A, 4A). This imaging also confirmed the presence of a hollow lumen within the organoids (Figure 3A, 4A).

**Figure 3.**
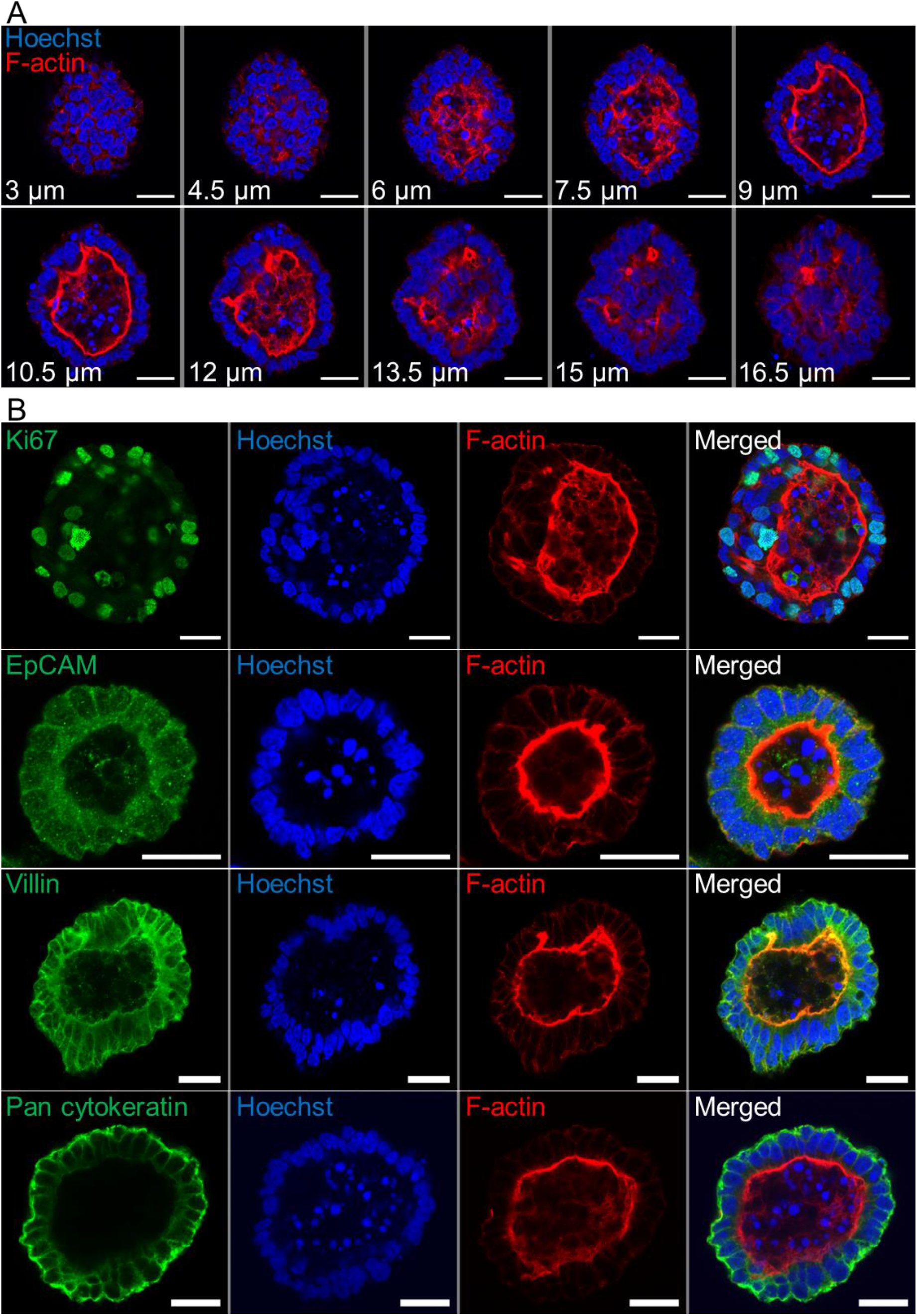
Characterisation of ovine abomasum organoids by immunofluorescence confocal microscopy. (A) Representative Z-stack images of an individual abomasum organoid with a closed luminal space and an internal F-actin-expressing brush border. Red = F-actin and blue = Hoechst (nuclei). (B) Representative images of abomasum organoids probed for either the cell proliferation marker Ki67, or the epithelial cell markers EpCAM, villin and pan-cytokeratin (all green). F-actin (red) and Hoechst (blue). Scale bars = 10 μm.

**Figure 4.**
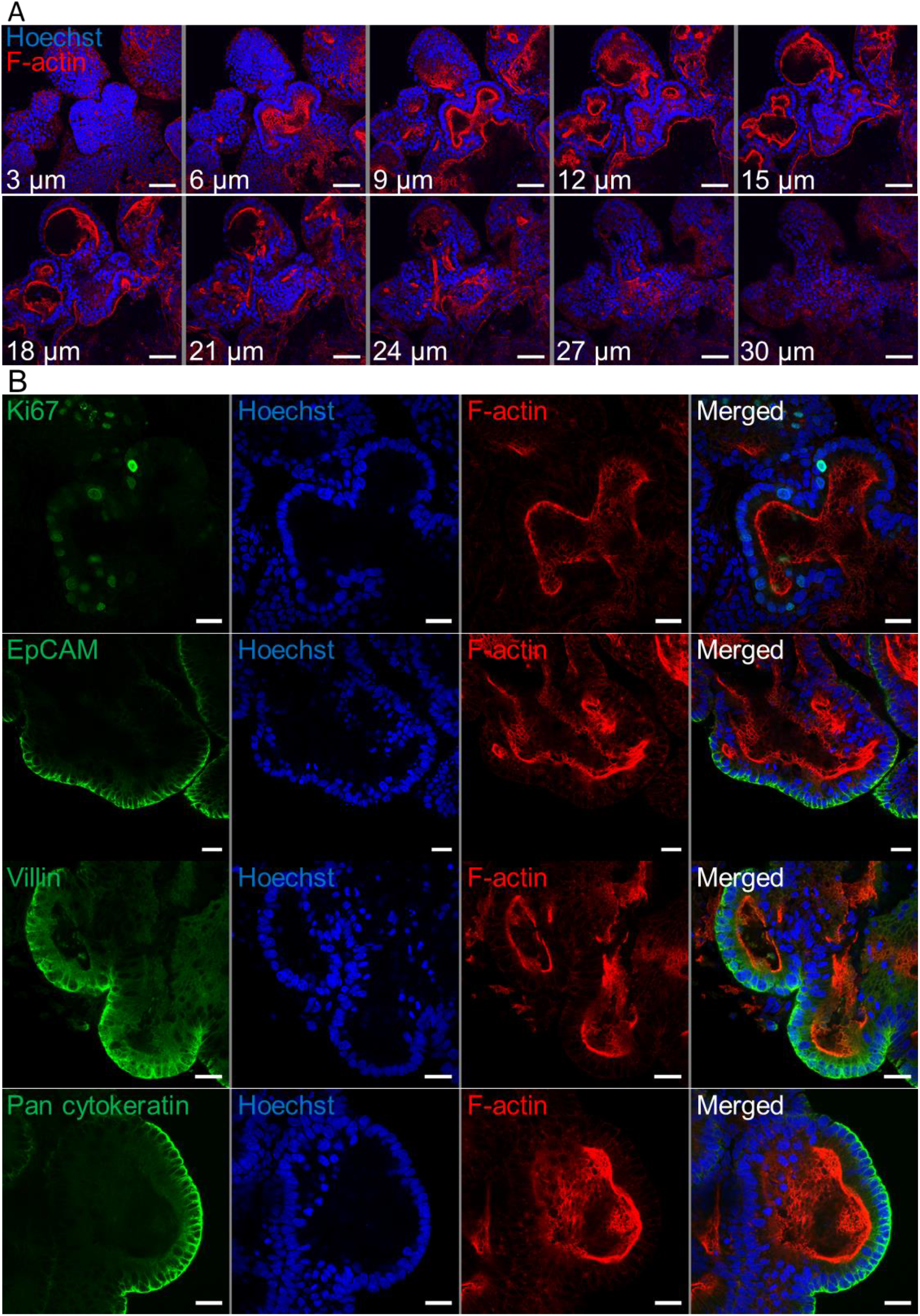
Characterisation of ovine ileum organoids by immunofluorescence confocal microscopy. (A) Representative Z-stack images of part of an individual branched ileum organoid with a closed luminal space and an internal F-actin-expressing brush border. Red = F-actin and blue = Hoechst (nuclei). (B) Representative images of abomasum organoids probed for either the cell proliferation marker Ki67, or the epithelial cell markers EpCAM, villin and pan-cytokeratin (all green). F-actin (red) and Hoechst (blue). Scale bars = 10 μm.

The proliferation marker Ki67 was detectable in both the abomasum and ileum organoids, indicating that cell division continued to take place after 7 days of *in vitro* culture (Figure 3B, 4B). The epithelial cell markers EpCAM (epithelium cell adherence molecule), villin (epithelium-specific actin-binding protein) and cytokeratin (epithelial cell cytoskeleton filament protein) were each detectable in abomasum and ileum organoids (Figure 3B, 4B), confirming the differentiation of stem cells into epithelium cell-containing organoids. Control samples of organoids probed with mouse and rabbit serum IgG did not label positive for any of the epithelial cell markers, confirming the specificity of the epithelial cell labelling (Figure S1, S2).

### 3.3 Transcriptional analysis of abomasum and ileum organoids and tissue

Gene expression profiles from: ovine ileum organoids (P0 – P4); ovine abomasum organoids (P0 – P4); ovine ileum tissue (n = 5) and ovine abomasum tissue (n = 5) were compared by RNA-seq analysis. The global gene expression profiles of the complete dataset, consisting of 20 individual samples, were initially compared by principal component analysis (PCA) (Figure 5). The PCA analysis resulted in four statistically significant clusters(with 95% confidence intervals), with each cluster representing a sample type (i.e. ileum organoids; abomasum organoids; ileum tissue; or abomasum tissue). This initial analysis demonstrates that replicate samples collected from either ileum tissue (n = 5) or abomasum tissue (n = 5) group by tissue type showing that the global transcript signature of ileum and abomasum tissue differ. Based on global gene expression profiles, organoids also grouped by the tissue type from which they derived, with ileum and abomasum organoids forming separate statistically significant clusters (95% confidence intervals) in the PCA analysis (Figure 5). Importantly, for both ileum and abomasum organoids, each passage (P0 – P4) is represented in each cluster, showing that there was no global change in the transcriptome profile following serial passage (Figure 5).

**Figure 5.**
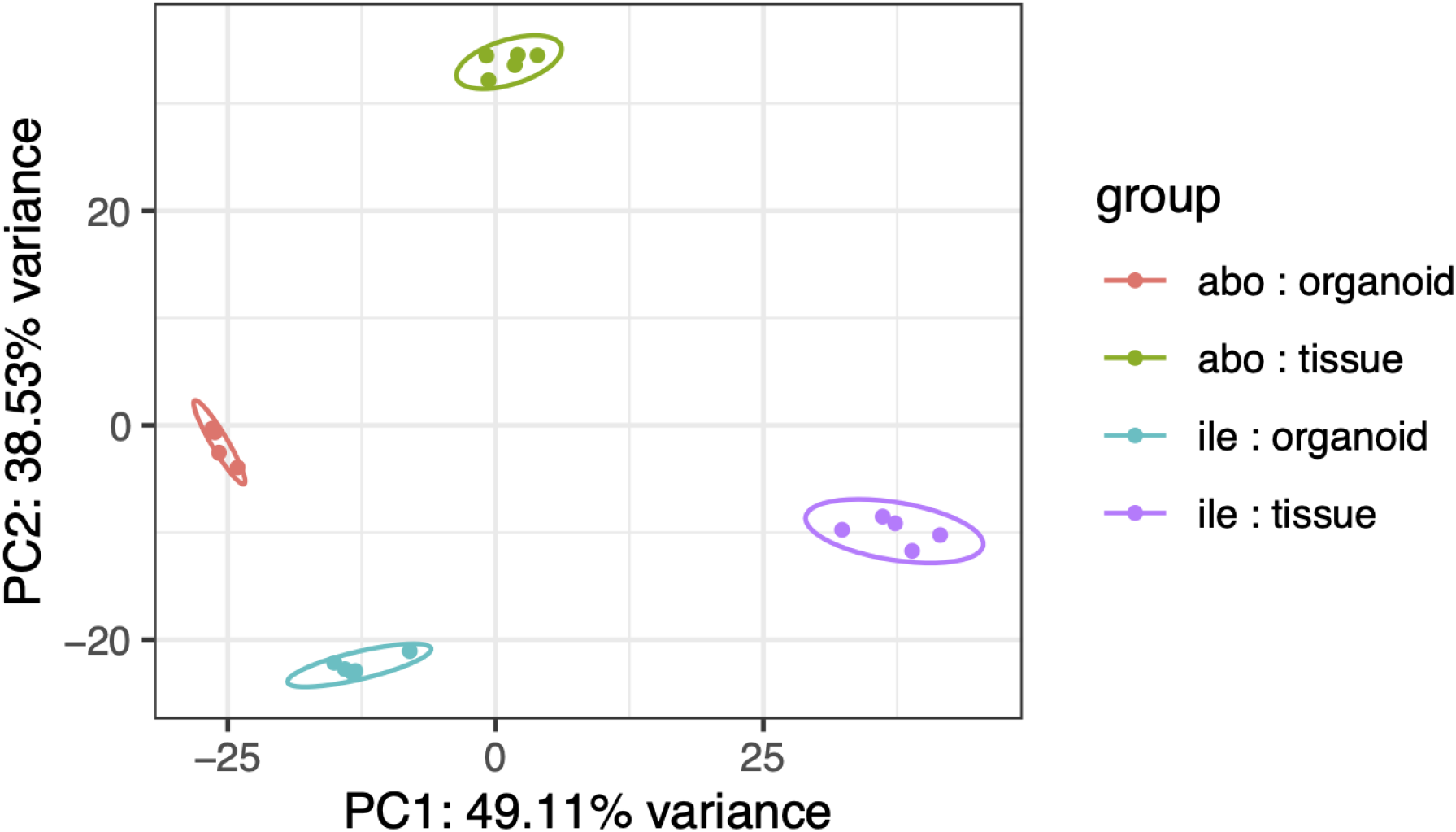
Principal component analysis (PCA) of RNA-seq expression of the top 500 most variant genes (of genes ranked by inter-sample variance) in ovine abomasum and ileum organoid and tissue samples. Sample type is indicated in the key and includes: abomasum organoid (red); abomasum tissue (green); ileum organoid (blue); ileum tissue (purple). Ellipses indicates 95% confidence intervals for each cluster.

The expression profiles of the top 40 most variable genes (of genes ranked by inter-sample variation) were compared from ileum and abomasum organoids from serial passages (P0 – P4) and ileum and abomasum tissue derived from five lambs (n = 5) (Figure 6). This analysis broadly identified three categories of genes, including genes with: i) abomasum (tissue and organoid) specific expression; ii) ileum (tissue and organoid) specific expression and iii) ileum and abomasum (tissue only) expression.

**Figure 6.**
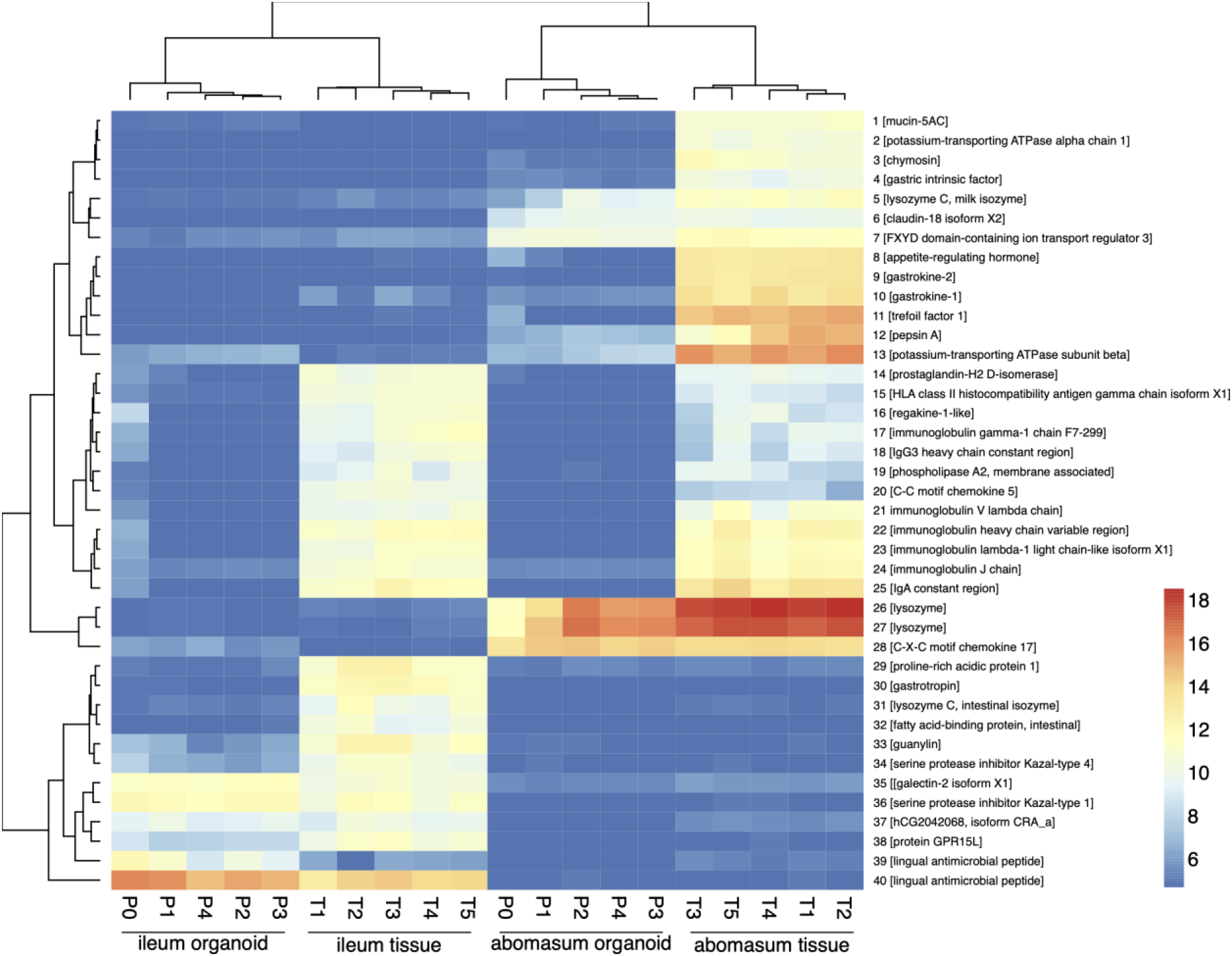
Heat map showing expression level of top 40 most variant genes (of genes ranked by inter-sample variance) from ileum (ile) and abomasum (abo) organoids from serial passages (P0 – P4) and ileum (ile) and abomasum (abo) tissue derived from five lambs (T1 – T5). Colours indicate level of expression from low (blue) to high (red). The dendrograms indicate similarity between samples. Details of genes included in the heat map, including ENSOART sequence identifiers, are shown in Supplemental File 1.

Based on gene expression profiles, genes that were highly expressed in abomasum tissue and abomasum organoids, but absent in all ileum samples, included genes of known gastric function, such as: *claudin-18*; *gastrokine*; *gastric lysozyme* and *pepsin*. Similarly, ileum specific genes were detected in both ileum tissue and ileum organoid samples, but absent from all abomasum samples included: *galectin*; *lingual antimicrobial peptide*; *guanylin* (a 15 amino acid peptide secreted from goblet cells). Interestingly, genes shared by ileum tissue and abomasum tissue, but largely absent from organoid cultures, were predominantly immune related genes (such as: C-C motif chemokine 5, regakine-1-like and various immunoglobulin chains) and likely reflect the presence of immune cells in ileal and abomasal mucosal tissue samples, which were not represented in ASC derived ileum and abomasum organoids. In summary, based on transcriptional profiles, abomasum and ileum organoids are broadly representative of the tissues they were derived from and appear to be transcriptionally stable over multiple passages.

### 3.4 Expression of cell- and tissue-specific genes in abomasum and ileum organoids and tissues

The ovine gastrointestinal transcriptomic database generated here was manually searched for genes that are representative of specific cell and tissue markers. A total of 151 genes were searched in this way and their expression in abomasum and ileum organoids and tissue was presented in heat maps. A number of cell junction markers were consistently expressed in both organoids and tissue, including genes encoding proteins related to tight junctions, gap junctions, adherens junctions and desmosomes (Figure S3).

We identified genes associated with particular epithelial cell subpopulations that were consistently expressed in abomasum and ileum organoids across multiple passages (P0-P4), as well as in ileum and abomasum tissue samples from five individual animals. These include numerous markers associated with stem cells, enterocytes, secretory and mucus-producing cells and Paneth cells (Figure 7). In particular, expression of the stem cell marker LGR5 was higher in both abomasum and ileum organoids compared to the respective tissue samples, indicating the presence of a relatively higher stem cell subpopulation in the organoids compared to tissues (Figure 6). Three enterocyte genes associated with ileum tissue were not detected in ileum organoids, namely *ALPI*, *APOA4* and *APOC3* (Figure 7). These enterocyte markers were not detected in abomasum organoids or abomasum tissue from any of the five individual animals. Expression of several genes associated with homeostasis in gastrointestinal cells was conserved between tissue samples and organoids, for both abomasum and ileum samples. This included *HES1*, *ADAM10*, *ADAM17*, *FGF20* and *SHH* (Figure 7).

**Figure 7.**
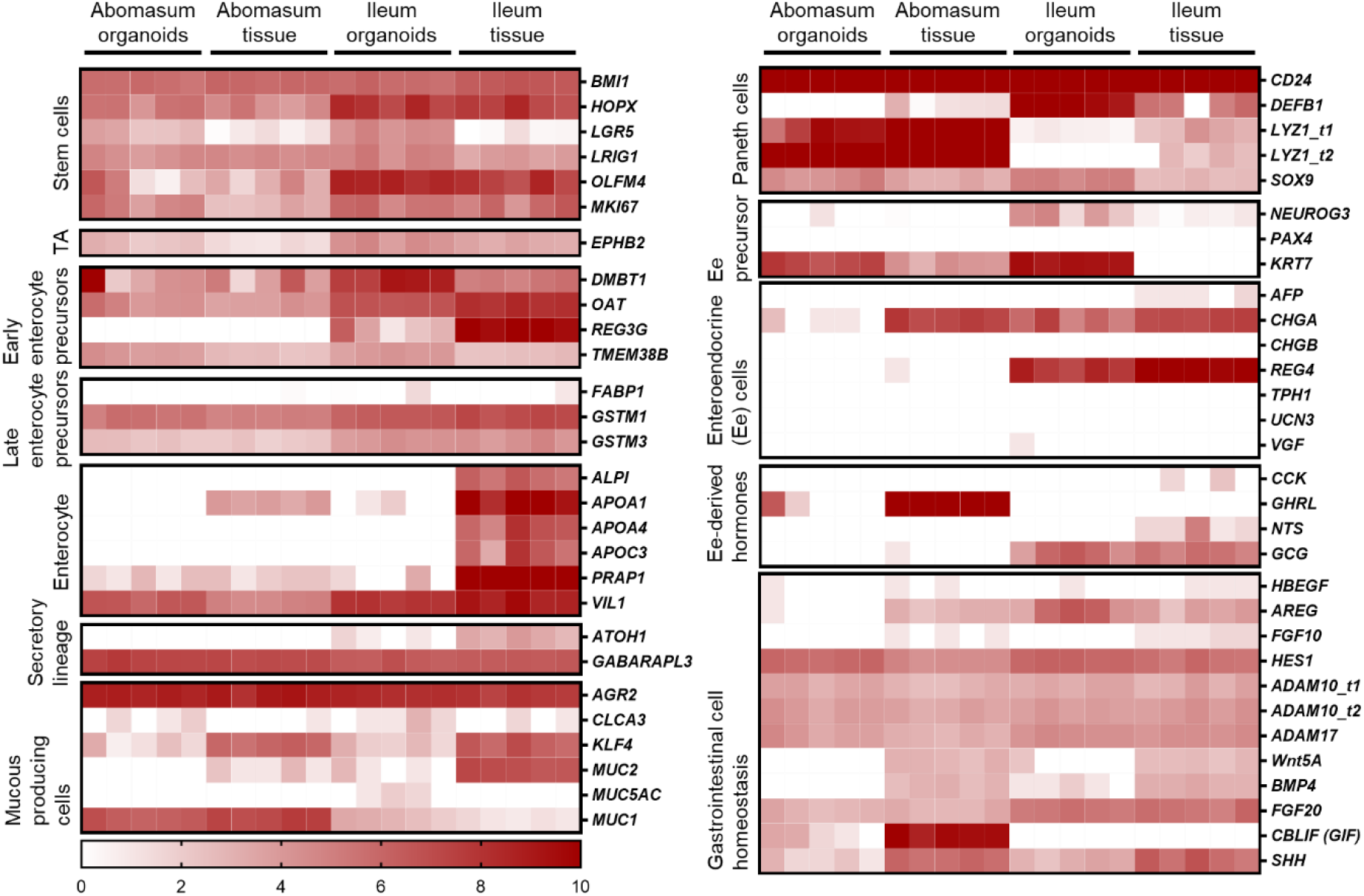
Heat map showing the expression of genes associated with gastrointestinal epithelia in abomasum and ileum tissue and organoids. RNA-seq analysis was performed to compare gene expression in abomasal and ileal tissue derived from five lambs and abomasum and ileum organoids across multiple passages. Squares from left to right under “abomasum tissue” and “ileum tissue” represent lambs T1-T5. Squares from left to right under abomasum organoids and ileum organoids represent passages P0-P4. Scale = log_2_ transcripts per million reads. Ee: enteroendocrine. Details of genes included in the heat map, including ENSOART sequence identifiers, are shown in Supplemental File 2.

A number of genes associated with specific epithelial cell subpopulations were differentially expressed in ileum and abomasum tissue. For example, the early enterocyte precursor-associated gene *REG3G*, the Paneth cell marker *DEFB1*, the enteroendocrine cell marker *REG4* and the enteroendocrine cell-derived hormone *GCG* were expressed in ileum tissue and not in abomasum tissue (Figure 7). These genes were also expressed in intestinal organoids and not abomasum organoids, indicating the conservation of tissue-specific differences in the cell subpopulations of the two different types of organoids.

Various genes were found to be specific for the abomasum, being expressed in both abomasal tissue and abomasum organoids but not in ileal tissue or ileum organoids. These included *PGA5, CCKBR* and *CBLIF (GIF)* (Figure 8). We also found that some genes specifically expressed in abomasal tissue were not expressed in abomasum organoids, including *SLC5A5, DUOX2, MCT9, PGC, ATP4A, AQP4,* and *HDC* (Figure 8).

**Figure 8.**
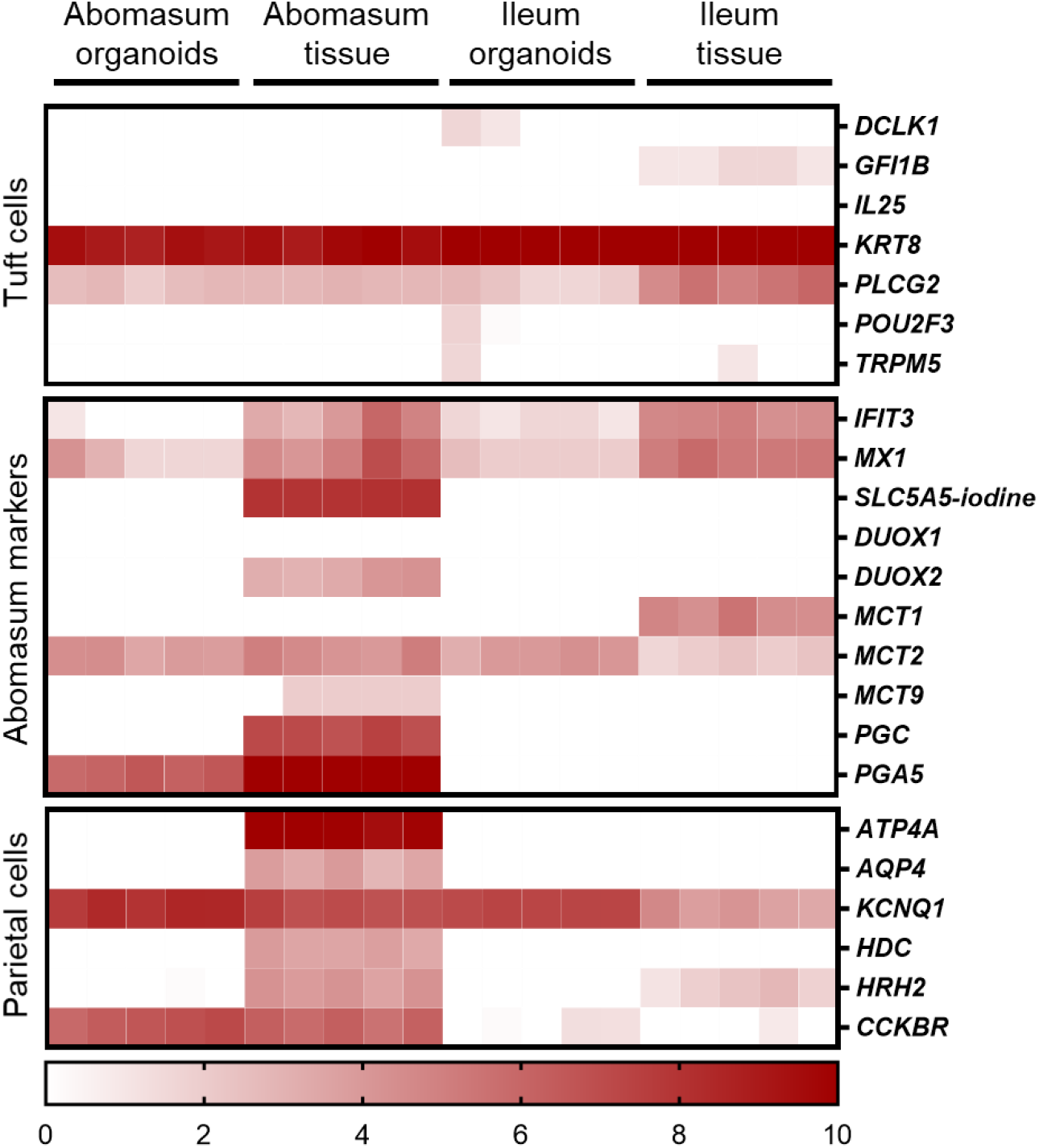
Heat map showing the expression of genes associated with gastric epithelia in abomasum and ileum tissue and organoids. RNA-seq analysis was performed to compare gene expression in abomasal and ileal tissue derived from five lambs and abomasum and ileum organoids across multiple passages. Squares from left to right under “abomasum tissue” and “ileum tissue” represent lambs T1-T5. Squares from left to right under abomasum organoids and ileum organoids represent passages P0-P4. Scale = log_2_ transcripts per million reads. Details of genes included in the heat map, including ENSOART sequence identifiers, are shown in Supplemental File 2.

The expression of immune-related genes, including toll-like receptors (TLRs), c-type lectin receptors (CLRs), chemokines, cytokines and antimicrobials were examined in abomasum and ileum tissue and organoids. The TLRs - *TLR3*, *TLR5* and *TLR6*, and CLR *Dectin-1* were expressed in abomasum and ileum organoids and their respective tissues (Figure S4). A number of chemokines were expressed in abomasum organoids and abomasal tissue, including *CXCL16*, *CCL20*, *CCL24* and *ACKR3*. Interestingly, the chemokine *CCL17* was up-regulated in abomasum and ileum organoids compared to the respective tissue samples (Figure S4). The expression of cytokine associated genes *IL18BP*, *IL27RA*, *IL4I1*, *IL13RA1* and *IFNGR1* was detected in abomasum and ileum organoids (Figure S4). Of note, the antimicrobial gene SBD2 was found to be highly expressed in ileum and ileal tissue, but was not expressed in either abomasum organoids nor abomasal tissue (Figure 6, Figure S4).

### 3.5 Organoid co-culture with the helminth *Teladorsagia circumcincta*

In order to use gastrointestinal organoids to study host-pathogen interactions *in vitro*, it is important to be able to challenge organoids with the pathogen-of-interest. Here, we co-cultured abomasum and ileum organoids with larvae of the important ruminant helminth parasite *T. circumcincta*. Infective, third stage larvae (L3) were ex-sheathed *in vitro* and labelled with the lipophilic dye PKH26. Labelled larvae were added directly to the well of a 24-well tissue culture plate containing abomasum or ileum organoids embedded in Matrigel and complete IntestiCult growth media. A number of *T. circumcincta* L3 penetrated the Matrigel, of which approximately 50% subsequently burrowed into central lumen of the organoids by 24 hours post-incubation, with some individual L3 invading the organoids as early as 2 hours. This indicated that it was possible to infect the organoids with the parasite in the correct orientation (i.e. with the parasite residing at the luminal surface of the organoid) without having to mechanically disrupt the organoids to allow access to the central lumen. *T. circumcincta* L3 were equally effective at infecting both abomasum and ileum organoids and motile larvae were still present after 14 days of co-culture. While we mainly observed abomasum organoids containing single larvae (Figure 9A), we found multiple larvae residing in the lumen of the larger ileum organoids (Figure 9B). Z-stack analysis on fixed samples showed worms were present within the lumen of the organoids and demonstrated L3 larvae burrowing directly through the epithelium of abomasum and ileum organoids to access the central lumen (Figure 9C).

**Figure 9.**
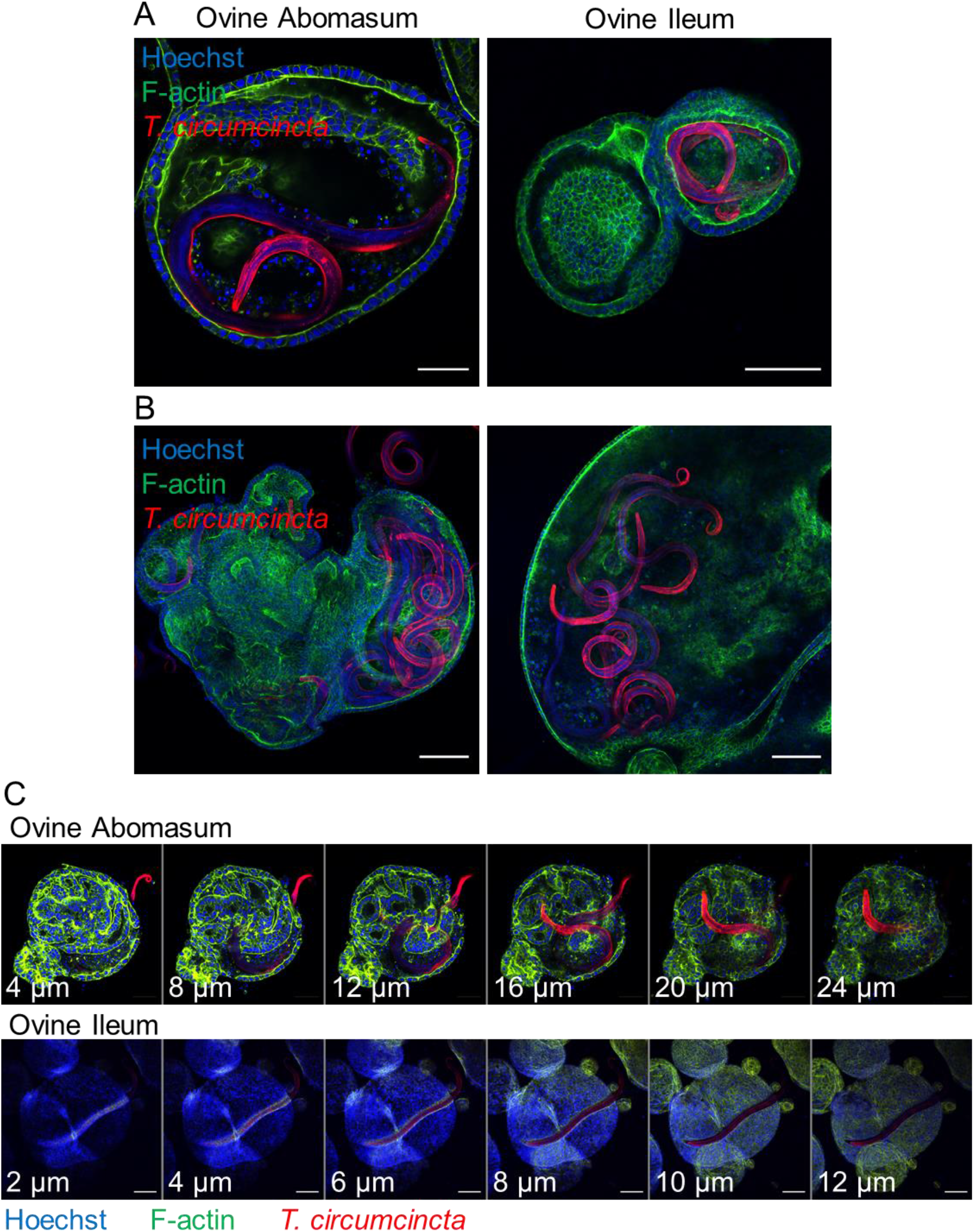
Ovine gastric and intestinal organoids modelling a helminth infection. (A) Representative images of ovine abomasum and ileum organoids challenged with the helminth parasite *Teladorsagia circumcincta*. Following 24 hours of co-culture, L3 stage T. circumcincta (red) are visible within the lumen of abomasum and ileum organoids. (B) Representative images of individual ileum organoids presenting an enlarged lumen containing multiple worms (red). (C) Representative Z-stack images showing L3 stage *T. circumcincta* (red) migrating through the epithelial layer in abomasum and ileum organoids. F-actin (green) and Hoescht (blue). Scale bars = 10 μm.

### 3.6 Generation of apical-out organoids and infection with *Salmonella typhimurium*

It is necessary to expose the apical surface of the organoid epithelia in order to have a working co-culture system for some pathogens. A recently published protocol (Co et al., 2019) described a method to invert the basal-out orientation of the abomasum and intestinal organoids. When the organoids were removed from Matrigel and incubated in 5 mM EDTA for 1 hour, the polarity of both the abomasum and intestinal organoids was reversed following 72 hours’ incubation in complete IntestiCult growth medium. F-actin staining of fixed organoid samples clearly highlighted the apical surface of the epithelium, which is initially internally located in basal-out abomasum and ileum organoids; however, after removing the extra cellular matrix from the organoids, the apical surface became positioned on the exterior surface of the organoids, with a microvilli brush edge apparent by confocal microscopy (Figure 10A, B).

**Figure 10.**
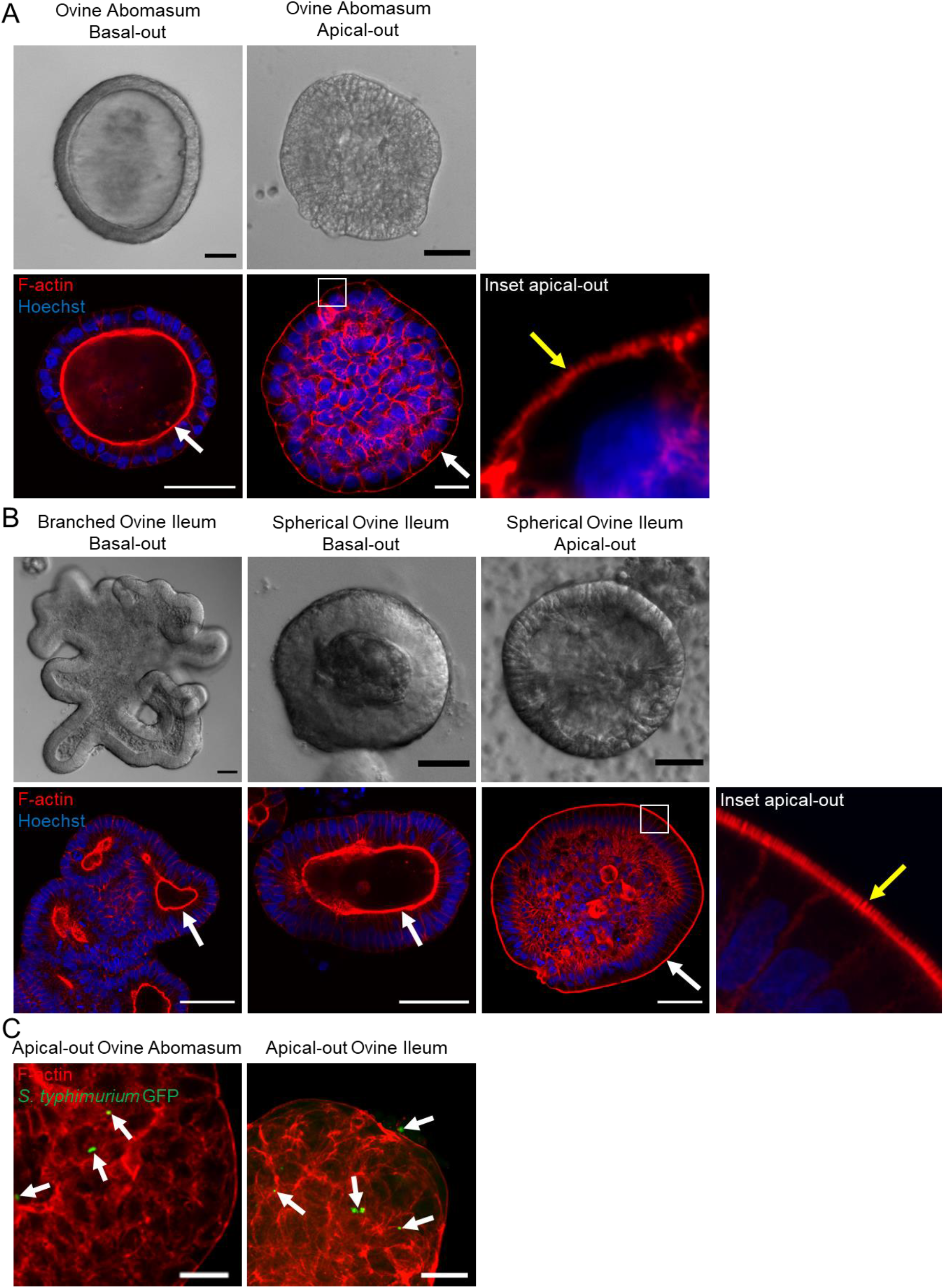
Reverse polarisation of ovine gastric and intestinal organoids for modelling host-pathogen interactions across the apical surface. Basal-out and apical-out abomasum (A) and ileum (B) organoids imaged by differential interference contrast (top) and confocal immunofluorescence microscopy (bottom). White arrows indicate the F-actin-expressing brush border associated with the apical surface of the epithelia. Yellow arrow in the inset panel indicates microvilli at the externally located brush border in apical-out organoids. (C) Cross sections of apical-out abomasum and ileum organoids imaged by confocal microscopy. GFP-expressing *Salmonella enterica* Typhimrium (green), indicated by white arrows, are detectable on the surface of and within epithelial cells. F-actin (red) and Hoechst (blue). Scale bars = 10 μm.

To demonstrate the utility of apical-out ovine gastric and intestinal organoids as an *in vitro* model for host-pathogen interactions, the apical-out organoids were exposed to the bacterial pathogen *Salmonella enterica* serovar Typhimurium, which is known to invade the epithelium via the apical surface (Finlay and Falkow, 1990). After 6 hours of organoid-bacteria co-culture freely suspended in complete IntestiCult growth medium, GFP-expressing *S.* Typhimurium were identifiable attached to the apical surface and within epithelial cells of the organoids by confocal microscopy. Although *S.* Typhimurium is an intestinal pathogen, here we observed GFP-expressing bacteria attached to both abomasum and ileum apical-out organoids (Figure 10C).

## 4. Discussion

Ruminants are key food-producing animals worldwide, providing a nutrient source to billions of people. Furthermore, dependency upon ruminants as a food source continues to increase in order to meet growing global dietary requirements. Gastrointestinal disease in ruminants is a major concern and accounts for significant economic losses and reduction in production efficiency. It is therefore important that ruminant health and welfare is improved through prevention and control of disease in order to meet ethical, economic and nutrient demands (Sargison, 2020).

An obvious challenge with studying gastrointestinal host-pathogen interactions *in vivo* is the internal nature of infections and the physical barriers associated with directly observing them. Therefore, a useful advancement for studying such infections is the development of a physiologically relevant *in vitro* model systems that allows experimental interrogation of host and pathogen interactions in fine detail. Stem cell-derived organoids have become a prominent feature of modern cell and tissue biology in recent years, representing *in vitro* cell cultures that retain structural and functional properties of the *in vivo* organ/tissue they represent (Clevers, 2016). To date, organoid cultivation has been achieved for numerous and diverse organs and tissues from different host species. In particular, organoids derived from gastrointestinal tissue have been generated for numerous livestock species, including cattle (Hamilton et al., 2018; Beaumont et al., 2021). However, the vast majority of these have been organoids representing the intestinal tract. Here, we demonstrated the ability to cultivate organoids from gastric and intestinal tissues of a small ruminant host and, to our knowledge, this is the first demonstration of organoids representing the gastric system of a ruminant.

Following the same protocol and using the same *in vitro* culture conditions, we report the ability to cultivate tissue-specific gastric and intestinal organoids from sheep. By comparing gene expression profiles between tissue and organoids, we found that when grown in identical conditions *in vitro*, stem cells from gastric glands developed into organoids that retained key characteristics associated with abomasum tissue. Stem cells from ileal crypts, on the other hand, developed into organoids which conserved important gene expression profiles associated with the ileum.

Ruminants, including cattle, sheep and goats are polygastric, in that they have a four-chambered gastric system. The fourth chamber, the abomasum, is most closely akin to the stomach of monogastric animals. An important differentiating characteristic between abomasum and ileum tissue is the expression of the digestive stomach enzyme pepsinogen in the abomasum (Mostofa et al., 1990). Another digestive protease associated with the abomasum in ruminants is lysozyme, which is highly expressed in this compartment (Stevens and Hume, 1998). Importantly, we found that both pepsinogen and lysozyme are expressed in abomasum organoids and not in ileum organoids. We also found evidence of parietal cells specifically present in abomasum organoids and not ileum organoids. This was indicated by the detection of *CCKBR* mRNA only in abomasum organoids and tissue following transcriptomic analysis. *CCKBR* is a cholecystokinin receptor expressed in the gastric and central nervous systems and more specifically it is associated with parietal cells in the stomach (Kulaksiz et al., 2000; Schmitz et al., 2001; Engevik et al., 2019). Conversely, we also identified genes whose expression was specific to ileum tissue that were also expressed in ileum organoids and not in abomasum organoids. For example, *REG4*, a marker of enteroendocrine cells (specifically enterochromaffin cells) in intestinal epithelia (Gehart et al., 2019) and *SBD2*, an antimicrobial sheep beta-defensin associated with the mucosal surface of small intestinal crypts (Meyerholz et al., 2004) were found to be specifically and abundantly expressed in ileum tissue and organoids and not abomasum. Collectively, these key differences in gene expression indicates that the two different types of organoid are tissue-specific and representative of the tissue from which the stem cells are derived. We also found that a number of genes used in previous studies as gastrointestinal epithelial markers (Hamilton et al., 2018) were not detected in our transcriptomic analysis of ileal or abomasal tissue from five individual animals, suggesting these genes are not reliable markers of gastrointestinal epithelia in sheep.

A necessary feature of an organoid cell line is the conservation of gene expression profiles across multiple passages. Transcriptomic analysis of abomasum and ileum organoid samples collected across five consecutive passages revealed that gene expression profiles were consistent. Further analysis of the expression of specific cell markers indicated that the diversity of epithelial cell types was also maintained across multiple passages. That the different organoid types maintain their tissue specificity and cell diversity, as well as the ability to cryopreserve them makes them a robust model that will ensure reproducibility across experiments, as well as reducing the reliance on deriving material from animals and thereby reducing the number of animals used in associated research.

To demonstrate the effectiveness of ovine gastric and intestinal organoids for modelling pathogen infections *in vitro*, we exposed abomasum and ileum organoids to different pathogens and showed they could invade them. It has been recognized that gastrointestinal organoids could represent useful *in vitro* models for studying helminth infections (Duque-Correa et al., 2020). However, to-date this has been limited to applying worm excretory and secretory products to organoids, or growing organoids from helminth-infected mice, as opposed to live host-parasite co-cultures (Eichenberger et al., 2018a, 2018b; Nusse et al., 2018; Luo et al., 2019; Duque-Correa et al., 2020). Here, we applied a very simple method of adding ex-sheathed *T. circumcincta* L3 directly to the growth media of organoids that were embedded in Matrigel. We found that after 24 hours, worms had burrowed through the Matrigel dome and into the lumen of individual organoids. We were also able to capture direct *T. circumcincta* invasion through the epithelium in both abomasum and ileum organoids. Furthermore, motile worms were observed at least 14 days following organoid invasion, demonstrating the potential to prolong parasite survival *in vitro* and to perform more long-term studies on the parasite compared to worms cultured under previous *in vitro* methods (*pers comms*).

Gastrointestinal pathogens that invade the epithelial mucosa commonly interact with the apical surface of epithelial cells. However, the innate polarity of mammalian gastrointestinal organoids grown in Matrigel is with the apical surface on the inside of the organoid. Various approaches have previously been used to expose pathogens to the apical surface of the epithelium, including microinjection directly into the lumen of the organoid, fragmentation of organoids and open-format 2D monolayers. A recent publication also demonstrated the ability to reverse the polarity of human ileum organoids by the removal of Matrigel and extracellular matrix proteins (Co et al., 2019). This has since been replicated in porcine ileum organoids (Beaumont et al., 2021) and here, we showed that ruminant ileum organoids can also have the polarity reversed following the same method. We also demonstrated that the polarity of gastric organoids can be reversed to an apical-out conformation. The ability to expose the apical surface of gastric and intestinal organoids to the culture supernatant facilitates direct interaction of the organoids with microbes, as we showed here by infecting apical-out organoids with *S.* Typhimurium. Since this method does not require the use of specialist equipment to administer pathogens into a central organoid lumen, this makes modelling host-pathogen infections *in vitro* significantly more practical.

In summary, the results from this study demonstrate the ability to isolate stem cells from gastric glands and crypts of the sheep abomasum and intestine, respectively and show that they differentiate into tissue-specific organoids when grown under identical conditions. The robustness of both gastric and intestinal organoids from sheep was demonstrated by showing that tissue-specific gene expression is maintained across multiple passages. Finally, both gastric and intestinal sheep organoids can be invaded by important bacterial and parasitic pathogens and they therefore represent a useful tool for modelling host-pathogen interactions.

## 6. Author Contributions

DS, DRGP, EAI and TMcN conceived the study. All authors designed the research. DS, DRGP, AB, KAH, MF, AFC and CC performed research. DS, DRGP and STGB analysed data. DS and DRGP wrote the paper with contributions from all authors. All authors read and approved the final manuscript.

## 7. Data availability statement

The datasets generated and analysed during the current study are fully compliant with the MINISEQE guidelines and are deposited in the publicly accessible NCBI Sequence Read Archive (SRA) Database under the project accession number PRJNA736945.

## 9. Acknowledgements

We would like to thank Leigh Andrews, Alison Morrison and Dave Bartley, Moredun Research Institute, UK, for their help and in the provision of parasite material and the Bioservices Unit, Moredun Research Institute, for expert care of the animals. We would also like to express our gratitude to Dr Prerna Vohra at the University of Edinburgh for their advice on the experimental infection of organoids with *S.* Typhimurium and for generation of the ST4/74 pFPV25.1 strain in a prior study.

## 11. Figure Legends

**Supplemental Figure 1.**
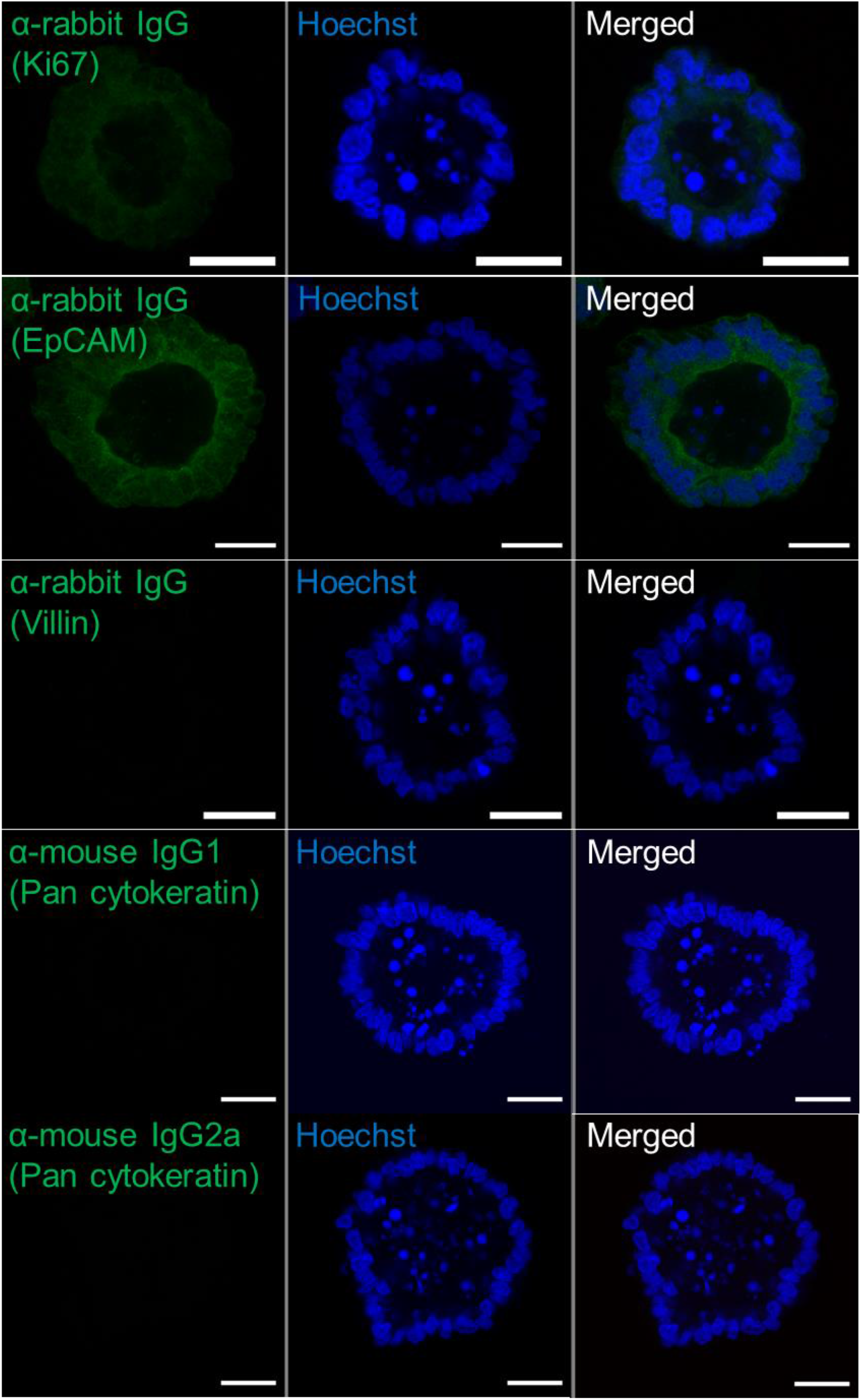
Negative controls for immunofluorescence antibody labelling in abomasum organoids. Representative confocal microscopy images of abomasum organoids probed with non-specific host IgG followed by indirect Alexa Fluor® 488-conjugated secondary antibody labelling. Marker name in green brackets indicates the antibody labelling control each organoid image represents. Scale bars = 10 μm.

**Supplemental Figure 2.**
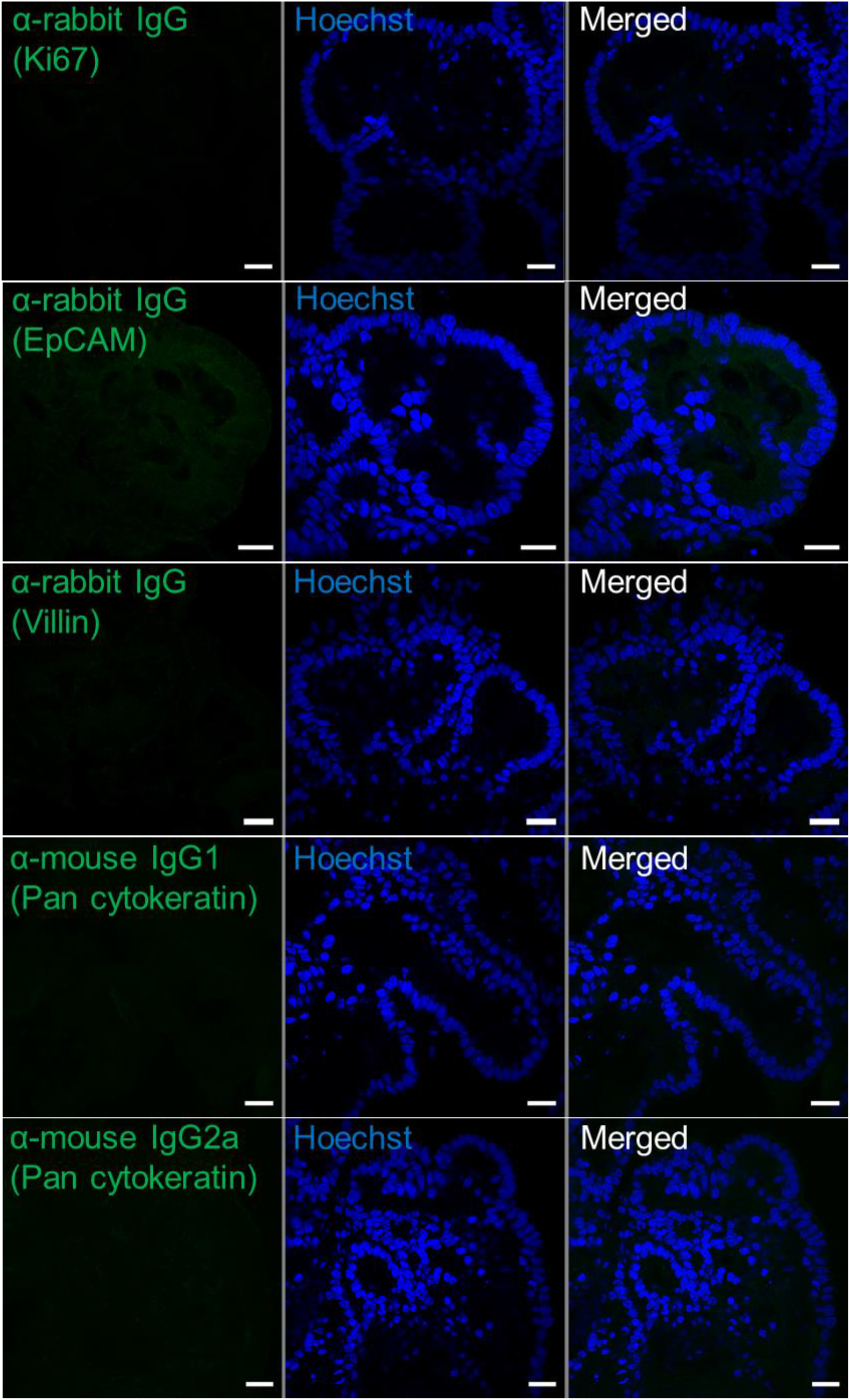
Negative controls for immunofluorescence antibody labelling in intestinal organoids. Representative confocal microscopy images of ileum organoids probed with non-specific host IgG followed by indirect Alexa Fluor® 488-conjugated secondary antibody labelling. Marker name in green brackets indicates the antibody labelling control each organoid image represents. Hoescht, blue. Scale bars = 10 μm.

**Supplemental Figure 3.**
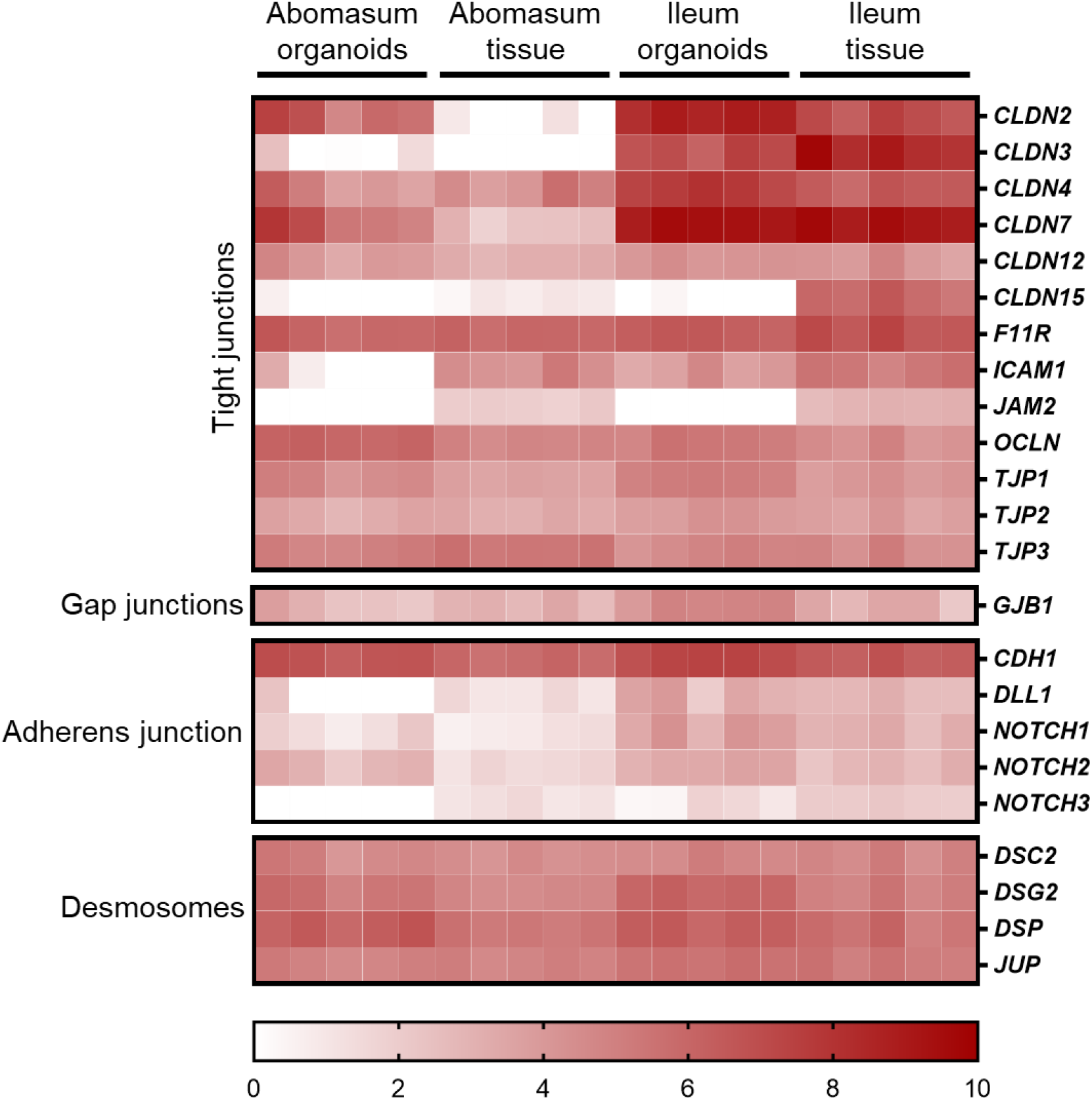
Heat map showing the expression of cell junction-related genes in abomasum and ileum tissue and organoids. RNA-seq analysis was performed to compare gene expression in abomasal and ileal tissue derived from five lambs and abomasum and ileum organoids across multiple passages. Squares from left to right under “abomasum tissue” and “ileum tissue” represent lambs T1-T5. Squares from left to right under abomasum organoids and ileum organoids represent passages P0-P4. Scale = log_2_ transcripts per million reads. Details of genes included in the heat map, including ENSOART sequence identifiers, are shown in Supplemental File 2.

**Supplemental Figure 4.**
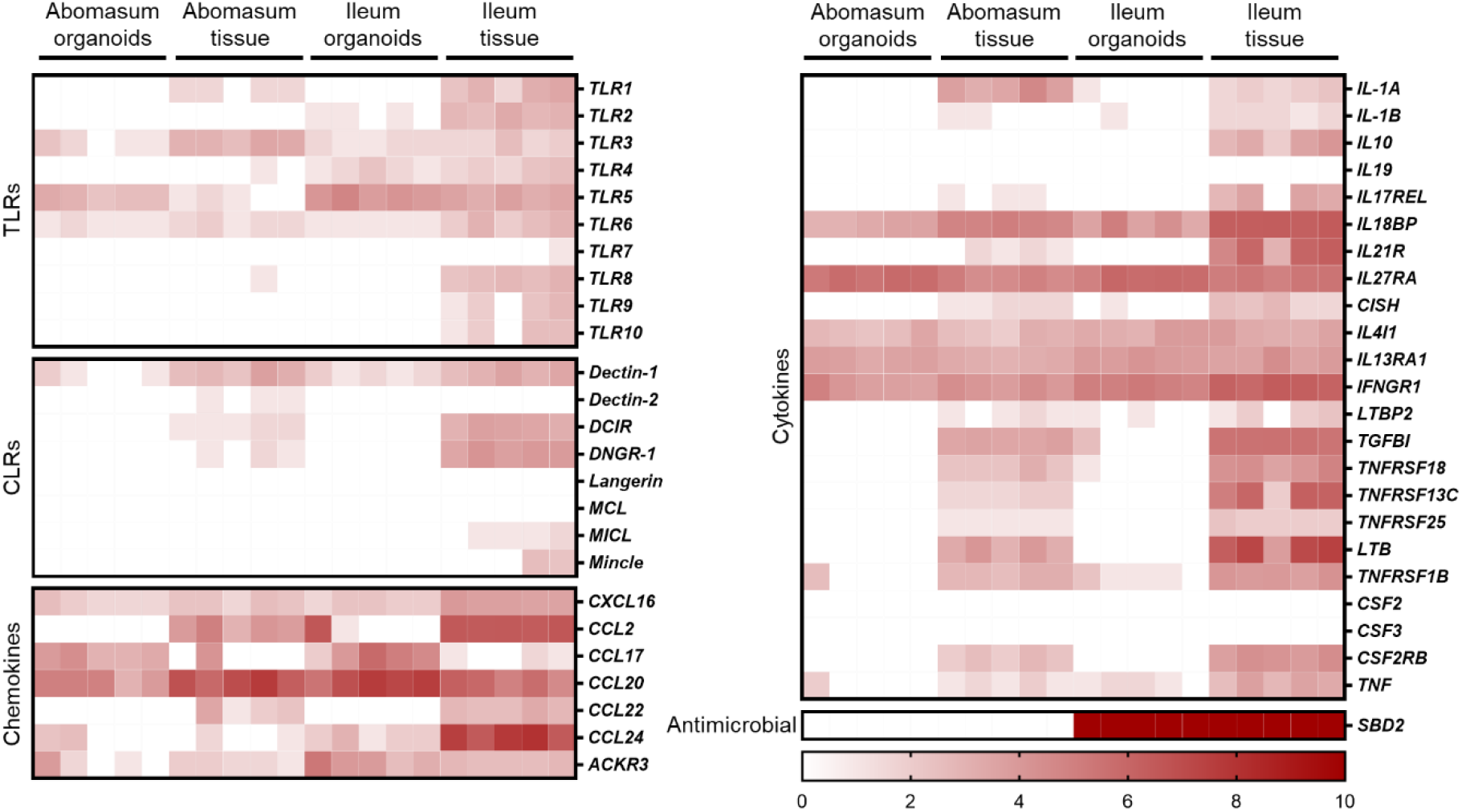
Heat map showing the detection of immune-related gene expression in abomasum and ileum tissue and organoids. RNA-seq analysis was performed to compare gene expression in abomasal and ileal tissue derived from five lambs and abomasum and ileum organoids across multiple passages. Squares from left to right under “abomasum tissue” and “ileum tissue” represent lambs T1-T5. Squares from left to right under abomasum organoids and ileum organoids represent passages P0-P4. Scale = log_2_ transcripts per million reads. TLRs: toll-like receptors. CLRs: C-type lectin receptors. Details of genes included in the heat map, including ENSOART sequence identifiers, are shown in Supplemental File 2.

